# Genomic identification, annotation, and comparative analysis of Vacuolar-type ATP synthase subunits in *Diaphorina citri*

**DOI:** 10.1101/2021.10.18.464890

**Authors:** Rebecca Grace, Crissy Massimino, Teresa D. Shippy, Will Tank, Prashant S. Hosmani, Mirella Flores-Gonzalez, Lukas A. Mueller, Wayne B. Hunter, Joshua B. Benoit, Susan J. Brown, Tom D’Elia, Surya Saha

## Abstract

Detailed annotation and comparative analysis were performed on the Asian citrus psyllid (ACP), *Diaphorina citri*, vacuolar-type ATP synthase (V-ATPase) to support the biological understanding and development of novel therapeutics to manage psyllid vectors. *D. citri* is a hemipteran insect that vectors the causative agent, the bacteria *Candidatus* Liberibacter asiaticus (*C*Las), of the citrus greening disease, Huanglongbing (HLB). Millions of citrus trees have been destroyed by citrus greening and every grove in Florida has been directly impacted. In eukaryotic organisms, V-ATPase is an abundant heterodimeric enzyme that serves the cell with essential compartment acidification through the active processes that transport protons across the membrane. Manual curation was completed on 15 putative *V-ATPase* genes in the *D. citri* genome. Comparative genomic analysis reveals that the *D. citri* V-ATPase subunits share domains and motifs with other insects, including the V-ATPase-A superfamily domain from the V-ATPase catalytic subunit A, which shares a 92% identity with *Acyrthosiphon pisum*. Phylogenetic analysis separates *D. citri* V-ATPase subunits into expected clades with orthologous sequences. Based on the results of annotation and comparative genomic analysis, RNAi therapies targeting *D. citri V-ATPase* genes, which have been successfully utilized in related hemipterans, are being pursued. Annotation of the *D. citri* genome is a critical step towards the development of directed-pest management that will lead to the reduced spread of the pathogens causing HLB throughout the citrus industry.

## Data Description

### Introduction

Vacuolar (H^+^)-ATP synthase (V-ATPase) is a highly conserved eukaryotic enzyme [1]. Originally identified in the vacuole membrane, V-ATPase has a critical function in the plasma membrane and endomembrane system of almost every cell [2,3]. V-ATPase works to regulate the acidity of organelles, such as vacuoles, the Golgi apparatus, and coated vesicles, by translocating protons across their membranes and powering secondary transport processes. Structurally, V-ATPase has a noncatalytic transmembrane domain, the V_0_ rotor, and a catalytic cytoplasmic domain, the V_1_ stator. V-ATPase hydrolyzes adenosine triphosphate (ATP) into adenosine diphosphate (ADP), thus acting opposite of the related F-ATPase [1]. In insects, 13 protein subunits are typically required to build a single V-ATPase [4]. The V_0_ domain consists of subunits a through e and V_1_ consists of subunits A through H [5]. There is also a critical accessory subunit S1 (Ac45) that helps assemble the enzyme [6].

### Context

In insects, high levels of V-ATPase are found in epithelial cells, and they are especially important in the digestive tract, helping to regulate nutrient uptake and solute transport [7]. Studies in several phyla, including insects, have demonstrated the lethality of silencing individual *V-ATPase* genes, making *V-ATPase* an attractive target for RNA interference (RNAi)-based pest control [1,4,7]. We have characterized the genes encoding V-ATPase subunits in *Diaphorina citri* (Hemiptera: Liviidae; NCBI:txid121845) as a step towards the development of future management strategies to reduce the psyllid vector of the causative agent of Huanglongbing (HLB), also known as citrus greening disease, the bacteria *Candidatus* Liberibacter asiaticus (*C*Las).

## Methods

Vacuolar ATP synthase insect orthologs from *Acyrthosiphon pisum* (pea aphid) were obtained from the KEGG database (RRID:SCR_012773). Additional ortholog subunits occurring in non-insect eukaryotes, like *Homo sapiens*, were obtained from HUGO Gene Nomenclature Committee (HGNC) (RRID:SCR_002827) and the non-redundant NCBI Reference Sequence database [8]. V-ATPase protein sequences were used to query the predicted protein set from the *D. citri* MCOT (Maker (RRID:SCR_005309), Cufflinks (RRID:SCR_014597), Oases (RRID:SCR_011896), and Trinity (RRID:SCR_013048)) transcriptome via BLASTp [9]. Reciprocal BLASTp analysis was performed to validate the *D. citri* MCOT significant hits using the NCBI non-redundant protein database [8]. *D. citri V-ATPase* genes were identified in the genome (version 1.91) by searching for the identified mapped MCOT models in the WebApollo (RRID:SCR_005321) system hosted at Boyce Thompson Institute. Multiple alignments of the predicted *D. citri* MCOT proteins, other gene model sequences, and insect *V-ATPase* orthologs were performed using the European Bioinformatics Institute MUSCLE alignment online tool (RRID:SCR_004727) [10]. Further analysis using RNA-seq reads, Illumina DNA-seq reads, StringTie models, and PacBio Iso-seq transcripts were used to manually annotate the final *V-ATPase* gene models. Manually annotated *V-ATPase* gene models were then integrated into the version 3.0 Official Gene Set (OGS). *V-ATPase* genes were verified in WebApollo through analysis using *de novo*-assembled transcripts, Iso-seq transcripts, Augustus models, Mikado transcriptome, SwissProt proteins, and SNAP prediction models. A list of annotated *D. citri* identifiers and a sampling of evidence that supports the annotated models are found in Table 1. A more detailed description of the annotation workflow is available (Figure 1) [11]. V-ATPase nomenclature is somewhat inconsistent in the literature and between species, therefore, we have used nomenclature standards reported in previous work on other Hemiptera [12,13].

**Table 1.** Evidence for gene annotations. MCOT transcriptome identifiers included, if applicable.

| Gene | Identifier | MCOT | *de novo*<br>transcriptome | Iso-seq | RNA-seq | Ortholog |
| --- | --- | --- | --- | --- | --- | --- |
| *V-ATPase a1* | Dcitr07g04330.1.1 | MCOT05340.0.CO | X | X | X | X |
| *V-ATPase a2* | Dcitr07g01670.1.1 | MCOT20572.0.CT | X | X | X | X |
| *V-ATPase b* | Dcitr12g08560.1.1 | - | X | X | X | X |
| *V-ATPase c* | Dcitr06g11110.1.1 | - | X | X | X | X |
| *V-ATPase d* | Dcitr04g06930.1.1 | - | - | X | X | X |
| *V-ATPase e* | Dcitr03g19730.1.1 | - | X | X | X | X |
| *V-ATPase A* | Dcitr06g09110.1.1 | MCOT04747.0.CC | - | X | X | X |
| *V-ATPase B* | Dcitr09g08730.1.1 | - | X | X | X | X |
| *V-ATPase C* | Dcitr02g01535.1.1 | - | X | X | X | X |
| *V-ATPase D* | Dcitr09g02030.1.1 | - | - | X | X | X |
| *V-ATPase E* | Dcitr04g09575.1.1 | - | - | X | X | X |
| *V-ATPase F* | Dcitr07g06920.1.1 | MCOT14638.0.CC | X | X | X | X |
| *V-ATPase G* | Dcitr11g08810.1.1 | MCOT22289.0.CT | - | X | X | X |
| *V-ATPase H*<br>(partial, N-terminus) | Dcitr00g06320.1.1 | - | - | - | - | - |
| *V-ATPase H*<br>(partial, C-terminus) | Dcitr01g01240.1.1 | MCOT00604.0.CT | - | X | X | X |
| *V-ATPase Ac45* | Dcitr09g09620.1.1 | MCOT16252.0.CC | X | X | X | X |

**Figure 1.**
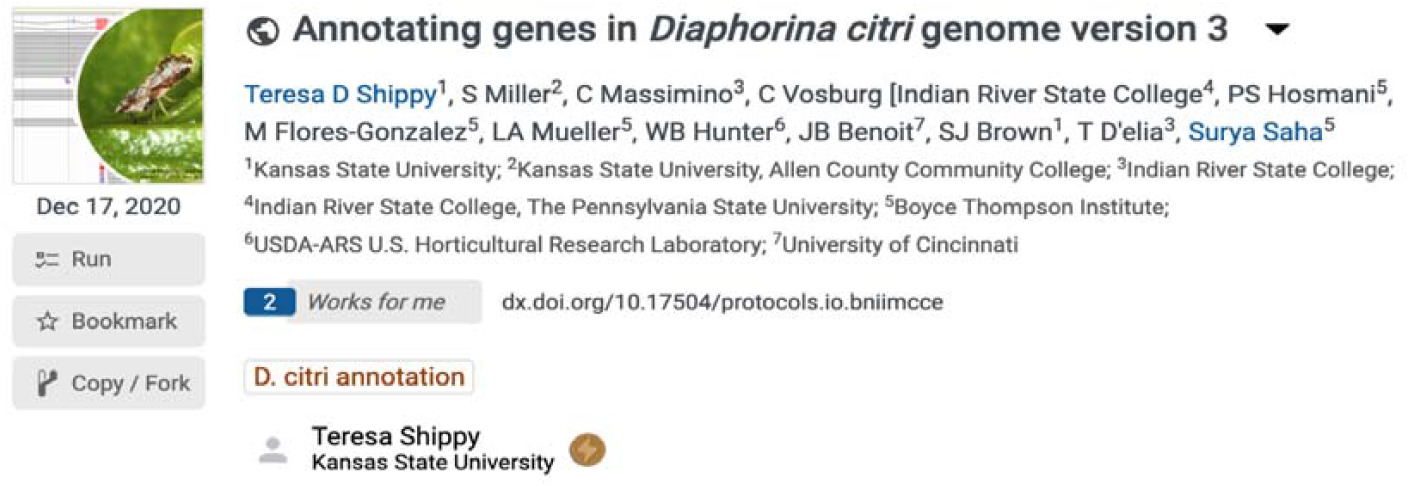
Protocols.io protocol for psyllid genome curation [11].

Reciprocal BLASTp of manually annotated v3.0 *V-ATPase* genes were performed at NCBI comparing the Insecta taxid. Insect orthologs from *Acyrthosiphon pisum* (pea aphid) [14,15], *Bemisia tabaci* (whitefly) [16], *Aedes aegypti* (yellow fever mosquito) [17], *Apis mellifera* (honeybee) [18], *Tribolium castaneum* (red flour beetle) [19], and *Drosophila melanogaster* (fruit fly) [20] were obtained by reciprocal BLASTp (RRID:SCR_004870) analysis of the non-redundant protein database at NCBI [8]. A neighbor-joining phylogenetic tree using the MUSCLE (RRID:SCR_011812) multiple sequence alignment with Poisson correction method and 1000 replicate bootstrap test was constructed using full-length protein sequences in MEGA version 7 (RRID:SCR_000667) for the transmembrane complex, the catalytic complex, and the accessory subunit Ac45, respectively (Figures 2-4) [21]. The sequence accession numbers used in these analyses can be found in Tables 2, 3, and 4. Comparative expression levels of *D. citri V-ATPases* throughout egg, nymph, and adult life stages in *D. citri* insects both exposed and not exposed to *C*Las were determined using RNA-seq data and the Citrus Greening Expression Network (CGEN) [9]. These gene expression levels were visualized using the pheatmap package in R (RRID:SCR_016418) [22,23]. Expression values for all samples discussed in this manuscript are visualized in Figures 5 and 6 and are reported as transcripts per million (TPM) in Table 5.

**Table 2.** *V-ATPase* transmembrane subunit (V_0_) BLAST table.

| Gene |  | *A. pisum* | *B. tabaci* | *T. castaneum* | *D. melanogaster* | *A. aegypti* | *A. mellifera* |
| --- | --- | --- | --- | --- | --- | --- | --- |
| *V-ATPase a1* | Accession | XP_029343960.1 | XP_018903507.1 | XP_008200952.1 | NP_650722.1 | XP_021706364.1 | XP_006565533.1 |
|  | Bit score | 1332 | 1312 | 1128 | 1151 | 1217 | 1166 |
|  | QC | 99% | 100% | 100% | 99% | 99% | 99% |
|  | Identity | 75.21% | 72.63% | 66.59% | 66.23% | 69.75% | 68.56% |
| *V-ATPase a2* | Accession | XP_008183003.1 | XP_018913655.1 | XP_008200806.1 | NP_733274.1 | XP_021693139.1 | XP_026298707.1 |
|  | Bit score | 1369 | 1334 | 1331 | 1286 | 1369 | 1392 |
|  | QC | 100% | 100% | 100% | 100% | 100% | 100% |
|  | Identity | 79.10% | 76.42% | 76.18% | 76.19% | 77.21% | 79.05% |
| *V-ATPase b* | Accession | NP_001155679.1 | XP_018909463.1 | NP_001161226.1 | NP_001247111.1 | XP_001662256.1 | XP_392599.1 |
|  | Bit score | 294 | 283 | 231 | 270 | 275 | 300 |
|  | QC | 99% | 99% | 98% | 99% | 99% | 99% |
|  | Identity | 73.79% | 73.30% | 63.05% | 72.95% | 74.4% | 76.33% |
| *V-ATPase c* | Accession | NP_001155531.1 | XP_018897791.1 | XP_967959.1 | NP_476801.1 | XP_001654757.1 | NP_001011570.1 |
|  | Bit score | 268 | 250 | 258 | 271 | 267 | 259 |
|  | QC | 98% | 99% | 98% | 99% | 98% | 98% |
|  | Identity | 92.11% | 86.93% | 89.54% | 92.16% | 92.76% | 89.47% |
| *V-ATPase d* | Accession | NP_001191854.1 | XP_018903442.1 | XP_974905.1 | NP_570080.1 | XP_001661299.1 | XP_393438.2 |
|  | Bit score | 656 | 669 | 671 | 661 | 677 | 679 |
|  | QC | 99% | 100% | 100% | 99% | 100% | 100% |
|  | Identity | 89.60% | 91.95% | 91.67% | 90.20% | 91.95% | 92.82% |
| *V-ATPase e* | Accession | XP_003242132.1 | XP_018909271.1 | XP_971898.1 | NP_001097499.1 | ABF18129.1 | XP_624787.1 |
|  | Bit score | 127 | 119 | 123 | 106 | 133 | 127 |
|  | QC | 96% | 96% | 94% | 96% | 100% | 94% |
|  | Identity | 69.51% | 70.73% | 72.50% | 58.54% | 70.59% | 71.25% |

Accession number, bit score, query coverage (QC), and identity results from protein BLAST analysis of annotated *D. citri* *V-ATPase*
transmembrane subunit genes to their putative orthologs.

**Table 3.** *V-ATPase* catalytic subunit (V_1_) BLAST table.

| Gene |  | *A. pisum* | *B. tabaci* | *T. castaneum* | *D. melanogaster* | *A. aegypti* | *A. mellifera* |
| --- | --- | --- | --- | --- | --- | --- | --- |
| *V-ATPase A* | Accession | XP_008179407.1 | XP_018897790.1 | XP_976188.1 | NP_001246015.1 | XP_021709029.1 | XP_016769524.1 |
|  | Bit score | 1176 | 1187 | 1167 | 1162 | 1168 | 1174 |
|  | QC | 99% | 99% | 99% | 99% | 100% | 99% |
|  | Identity | 91.84% | 92.47% | 90.21% | 90.18% | 90.57% | 90.72% |
| *V-ATPase B* | Accession | XP_003246082.1 | XP_018896879.1 | XP_967844.1 | NP_001163597.1 | XP_001651458.1 | XP_624112.1 |
|  | Bit score | 966 | 976 | 966 | 966 | 968 | 955 |
|  | QC | 98% | 99% | 98% | 98% | 98% | 98% |
|  | Identity | 93.88% | 94.33% | 93.48% | 93.88% | 94.09% | 92.87% |
| *V-ATPase C* | Accession | XP_001946227.1 | XP_018915661.1 | XP_008195426.1 | NP_477266.1 | XP_021695404.1 | XP_006562159.1 |
|  | Bit score | 702 | 653 | 671 | 645 | 671 | 648 |
|  | QC | 99% | 99% | 98% | 98% | 99% | 99% |
|  | Identity | 87.05% | 82.81% | 82.20% | 79.11% | 81.77% | 79.95% |
| *V-ATPase D* | Accession | NP_001119691.1 | XP_018904914.1 | XP_975872.1 | NP_651987.1 | XP_001660426.1 | XP_394769.2 |
|  | Bit score | 420 | 423 | 426 | 408 | 401 | 404 |
|  | QC | 100% | 100% | 100% | 100% | 100% | 100% |
|  | Identity | 87.24% | 86.42% | 86.53% | 82.52% | 80.89% | 86.94% |
| *V-ATPase E* | Accession | NP_001155650.1 | XP_018901912.1 | XP_970621.1 | NP_001287182.1 | XP_001655825.1 | XP_625098.1 |
|  | Bit score | 301 | 336 | 341 | 325 | 335 | 336 |
|  | QC | 99% | 100% | 100% | 100% | 100% | 100% |
|  | Identity | 73.21% | 71.68% | 73.01% | 71.24% | 70.80% | 73.01% |
| *V-ATPase F* | Accession | NP_001119690.1 | XP_018905603.1 | XP_975016.1 | NP_476969.1 | XP_001655376.1 | XP_624852.1 |
|  | Bit score | 223 | 227 | 216 | 217 | 216 | 230 |
|  | QC | 98% | 98% | 98% | 98% | 100% | 98% |
|  | Identity | 85.25% | 88.52% | 82.79% | 81.15% | 82.26% | 88.52% |
| *V-ATPase G* | Accession | NP_001119628.1 | XP_018908074.1 | XP_973974.1 | NP_001287407.1 | XP_001652605.1 | XP_624346.1 |
|  | Bit score | 184 | 188 | 173 | 154 | 171 | 180 |
|  | QC | 99% | 100% | 98% | 96% | 96% | 98% |
|  | Identity | 80.34% | 81.51% | 74.14% | 65.79% | 73.68% | 76.72% |
| *V-ATPase H*<br>(partial, N-terminus) | Accession | XP_001949116.3 | XP_018904009.1 | NP_001280516.1 | NP_001260510.1 | XP_001652018.1 | XP_003251675.1 |
|  | Bit score | 132 | 162 | 132 | 110 | 130 | 142 |
|  | QC | 90% | 99% | 99% | 84% | 91% | 94% |
|  | Identity | 57.69% | 68.42% | 57.26% | 56.70% | 58.10% | 64.22% |
| *V-ATPase H*<br>(partial, C-terminus) | Accession | XP_001949116.3 | XP_018904009.1 | NP_001280516.1 | NP_001260510.1 | XP_001652018.1 | XP_003251675.1 |
|  | Bit score | 530 | 506 | 513 | 493 | 498 | 490 |
|  | QC | 100% | 99% | 100% | 100% | 100% | 99% |
|  | Identity | 83.95% | 82.33% | 83.28% | 77.59% | 78.93% | 82.15% |

Accession number, bit score, query coverage (QC), and identity results from protein BLAST analysis of annotated *D. citri* *V-ATPase* catalytic subunit genes to their putative orthologs.

**Table 4.** *V-ATPase Ac45*, accessory subunit, BLAST table.

| Gene |  | *A. pisum* | *B. tabaci* | *T. castaneum* | *D. melanogaster* | *A. aegypti* | *A. mellifera* |
| --- | --- | --- | --- | --- | --- | --- | --- |
| *Ac45* | Accession | NP_001162140.1 | XP_018899028.1 | XP_974187.2 | NP_610470.1 | XP_001658652.1 | XP_001121483.3 |
|  | Bit score | 136 | 177 | 114 | 77.4 | 92.4 | 88.6 |
|  | QC | 83% | 97% | 100% | 96% | 95% | 93% |
|  | Identity | 30.87% | 33.49% | 27.03% | 24.94% | 23.81% | 26.35% |

Accession number, bit score, query coverage (QC), and identity results from protein BLAST analysis of annotated *D. citri* *V-ATPase*
*Ac45* gene to its putative orthologs.

**Table 5.**
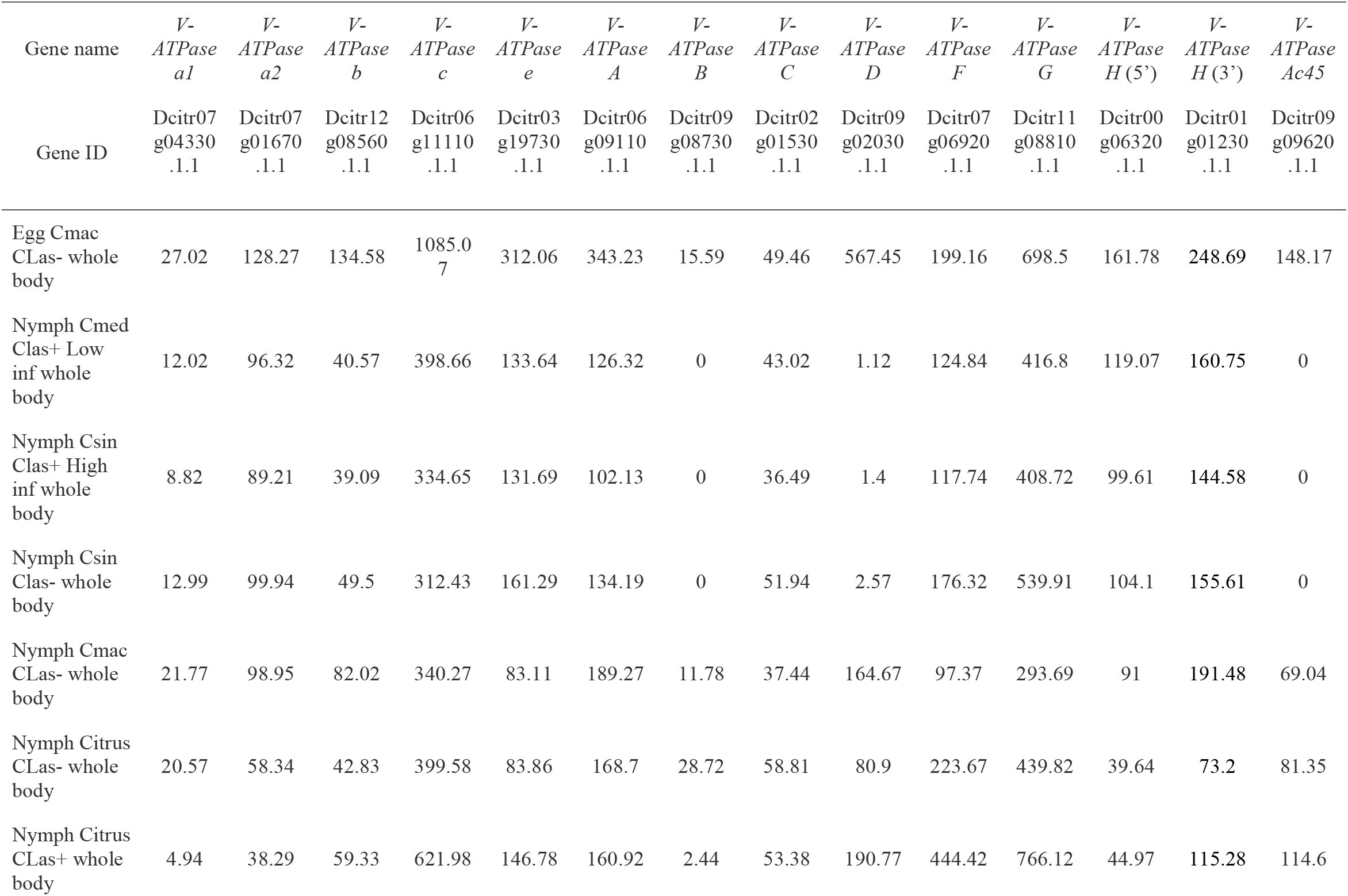
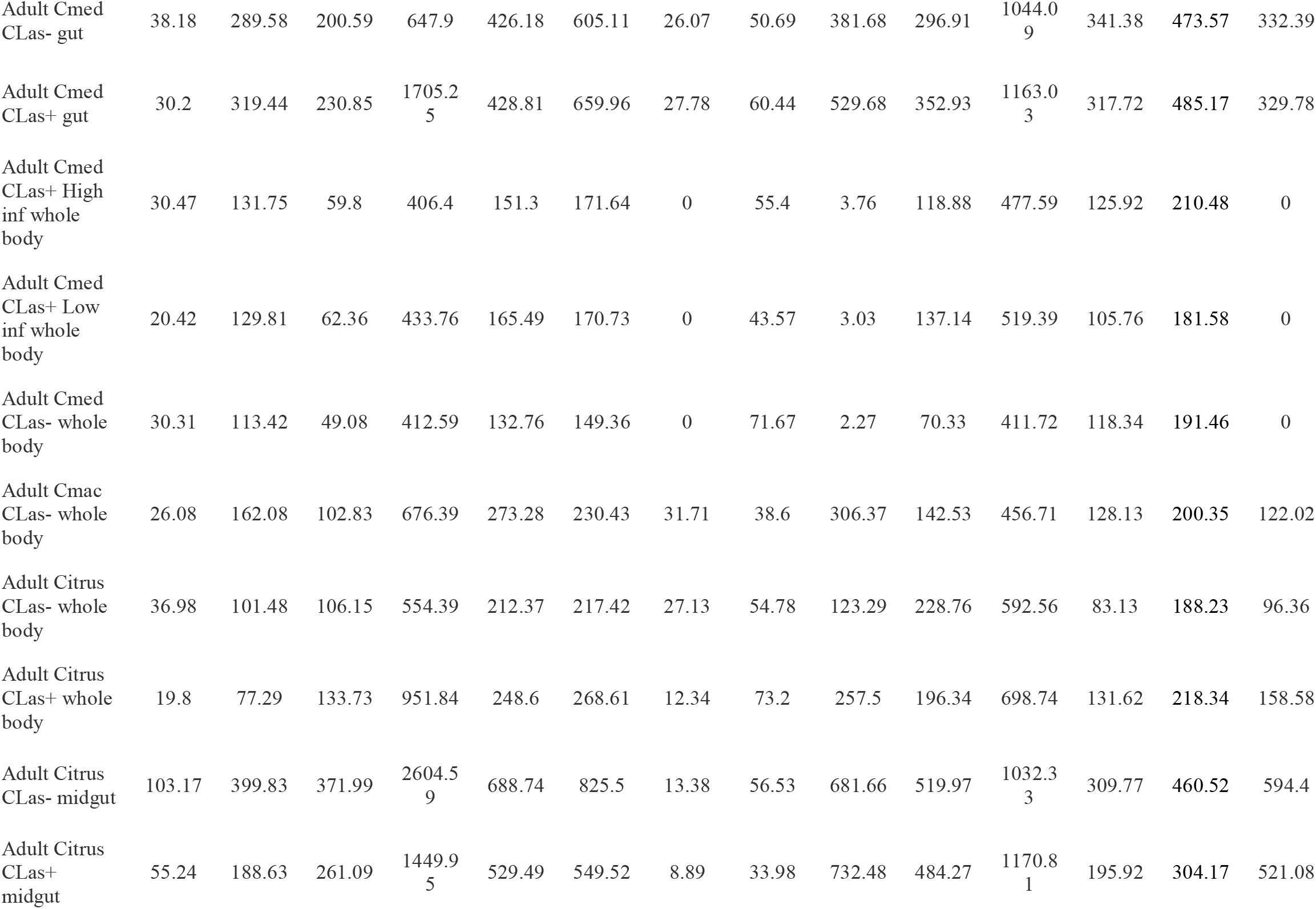
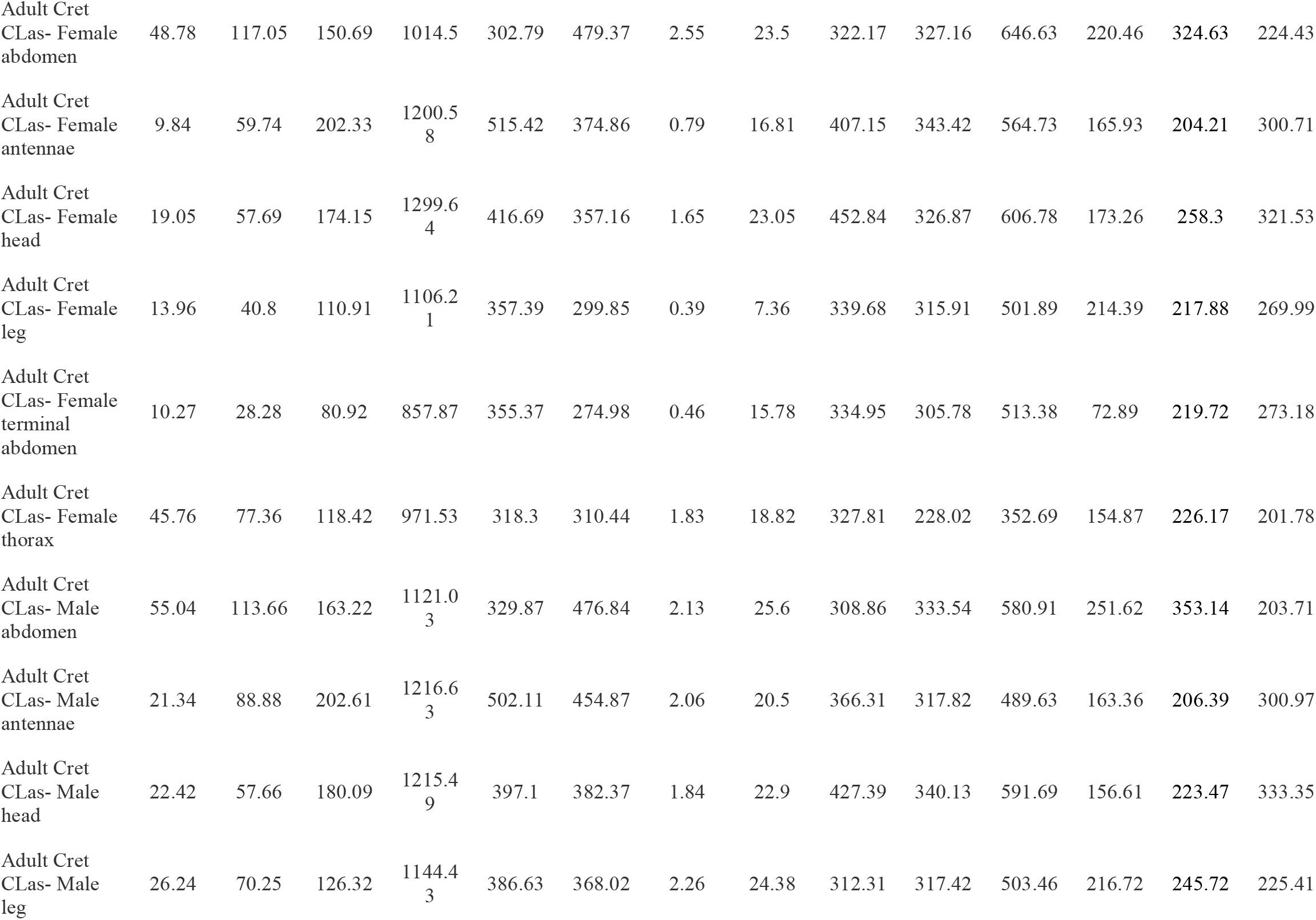
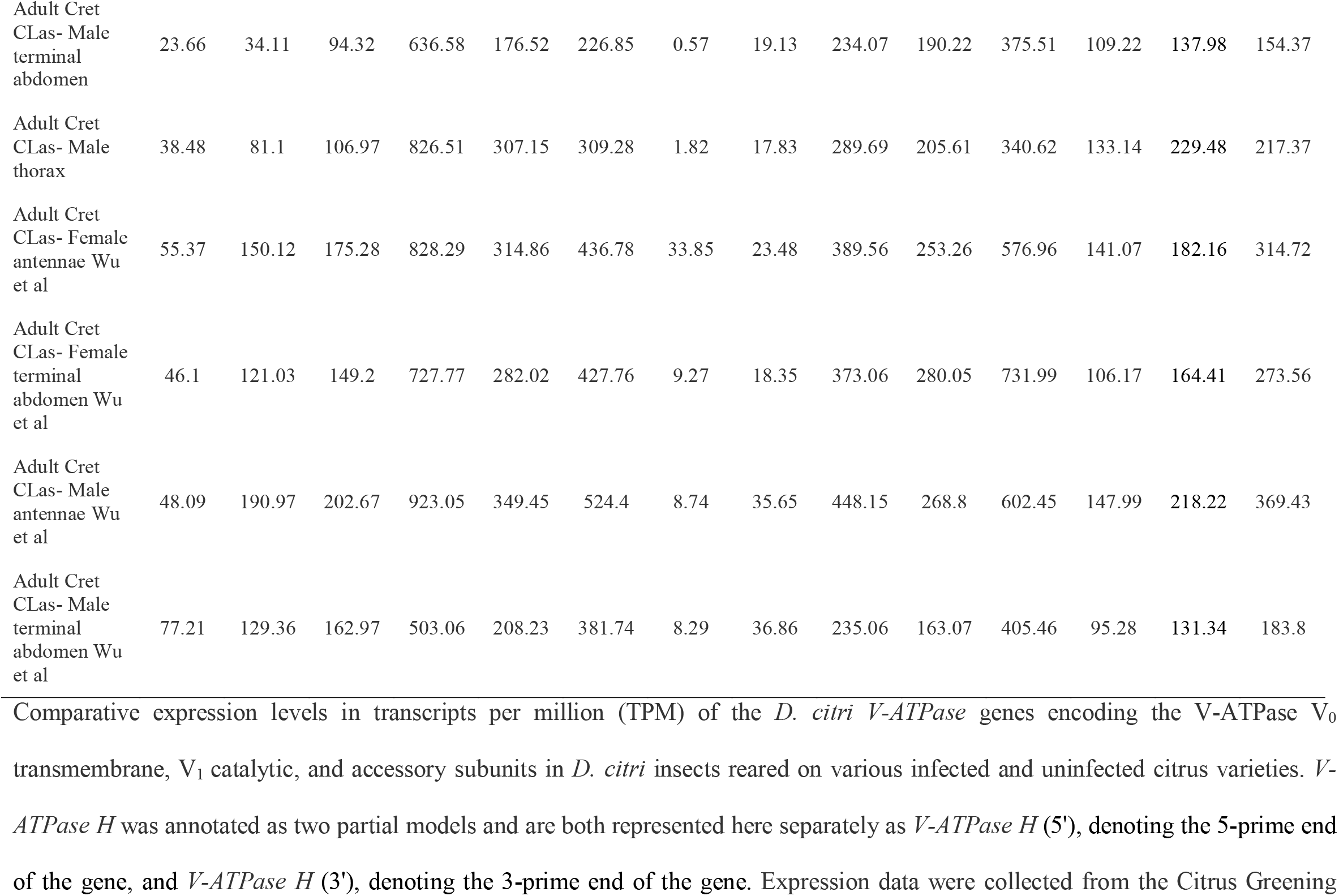
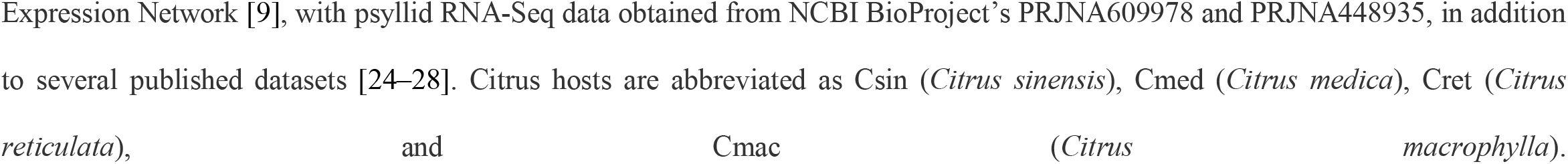
Expression values listed as TPM, visualized in Figures 5 and 6.

**Figure 2.**
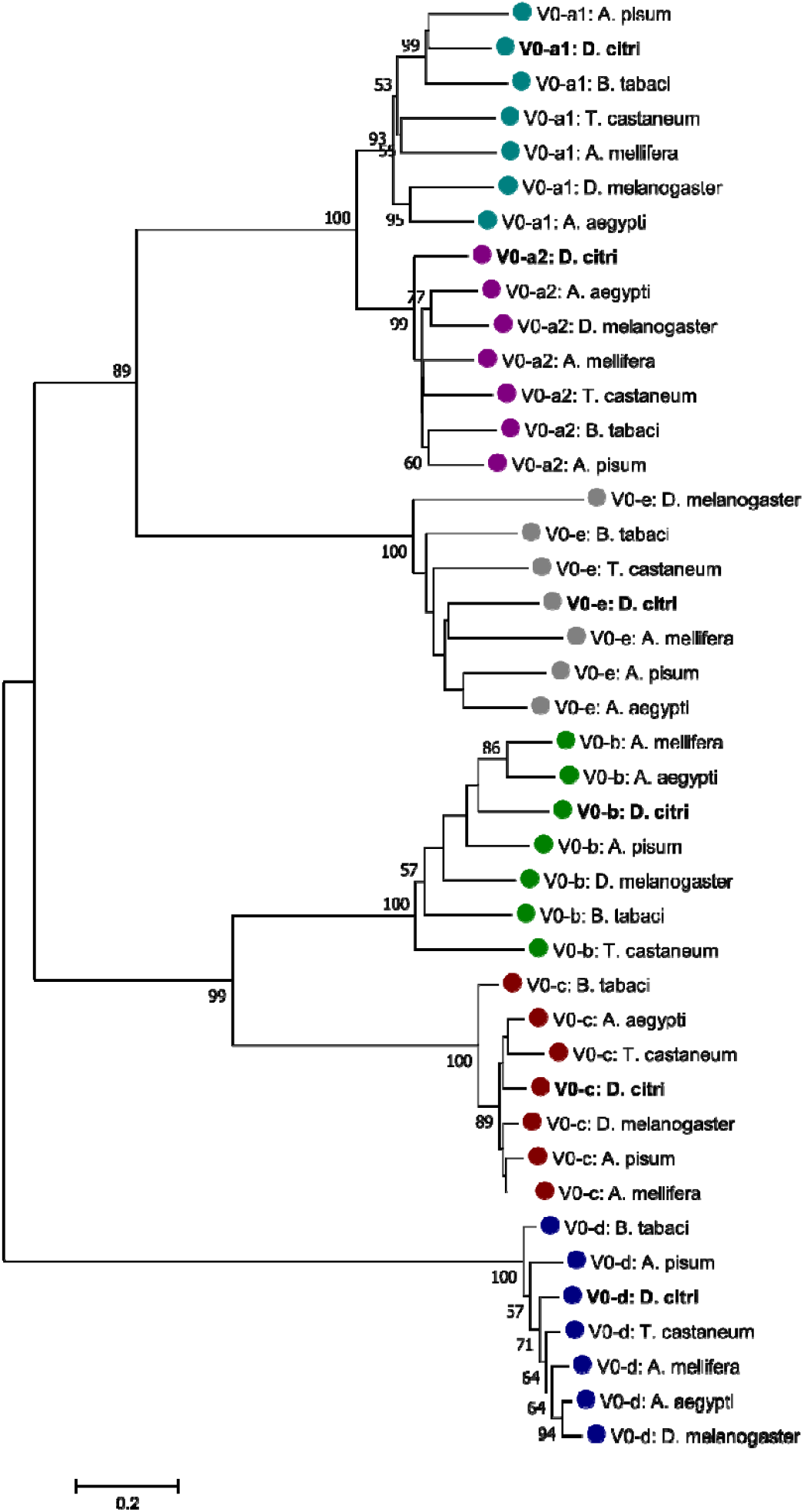
Phylogenetic analysis of V-ATPase V_0_, transmembrane domain, subunits a-e. The tree was constructed with MEGA7 software [21] using MUSCLE for alignment of amino acid sequences, followed by neighbor-joining analysis with 1000 bootstrap replications. Value greater than 50 are shown at nodes. *D. citri is* marked in bold and color-coding indicates specific V_0_ subunit groups. NCBI accession numbers are shown in Table 2.

**Figure 3.**
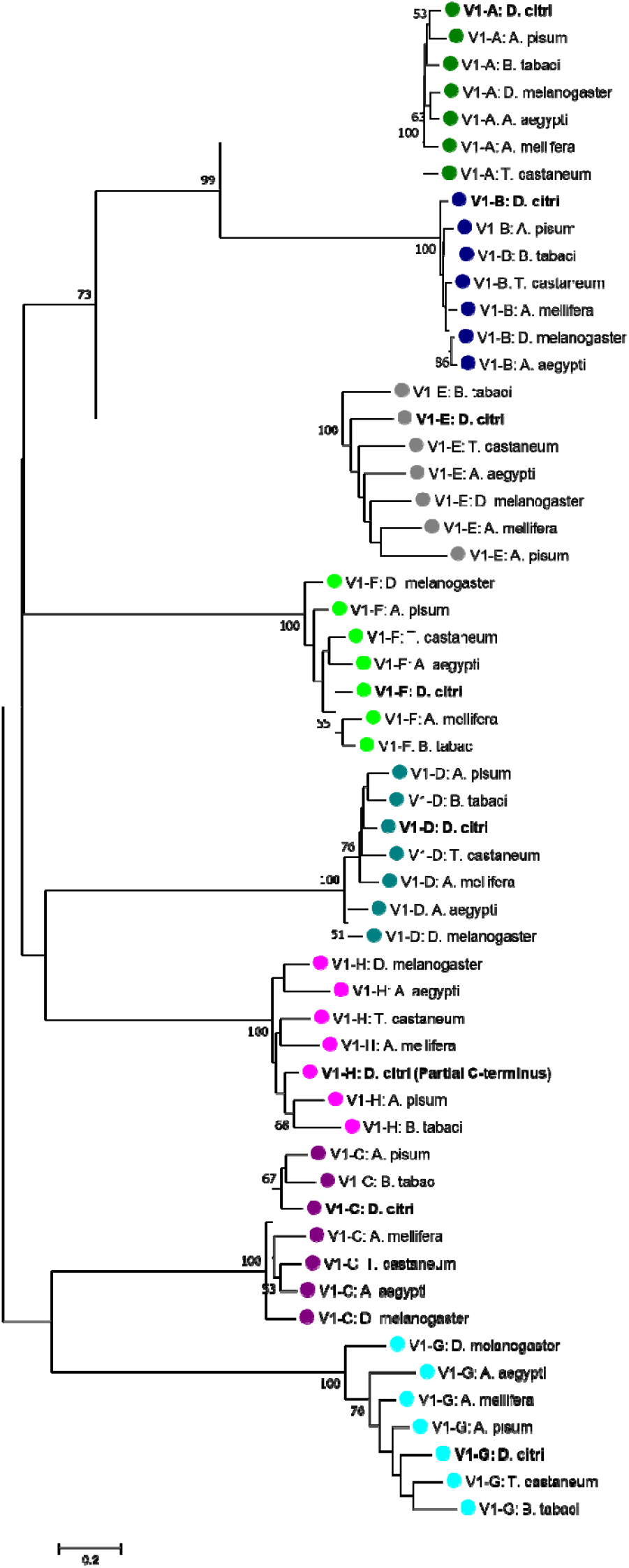
Phylogenetic analysis of V-ATPase V_1_, catalytic domain, subunits A-G. The tree was constructed with MEGA7 [21] software using MUSCLE for alignment of amino acid sequences, followed by neighbor-joining analysis with 1000 bootstrap replications. Values greater than 50 are shown at nodes. *D. citri is* marked in bold and color-coding indicates specific V_1_ subunit groups. *D. citri V-ATPase H* (V_1_-H) was annotated as two partial gene models, therefore, only the C-terminus, partial amino acid sequence of V_1_-H was included in this analysis. NCBI accession numbers are shown in Table 3.

**Figure 4.**
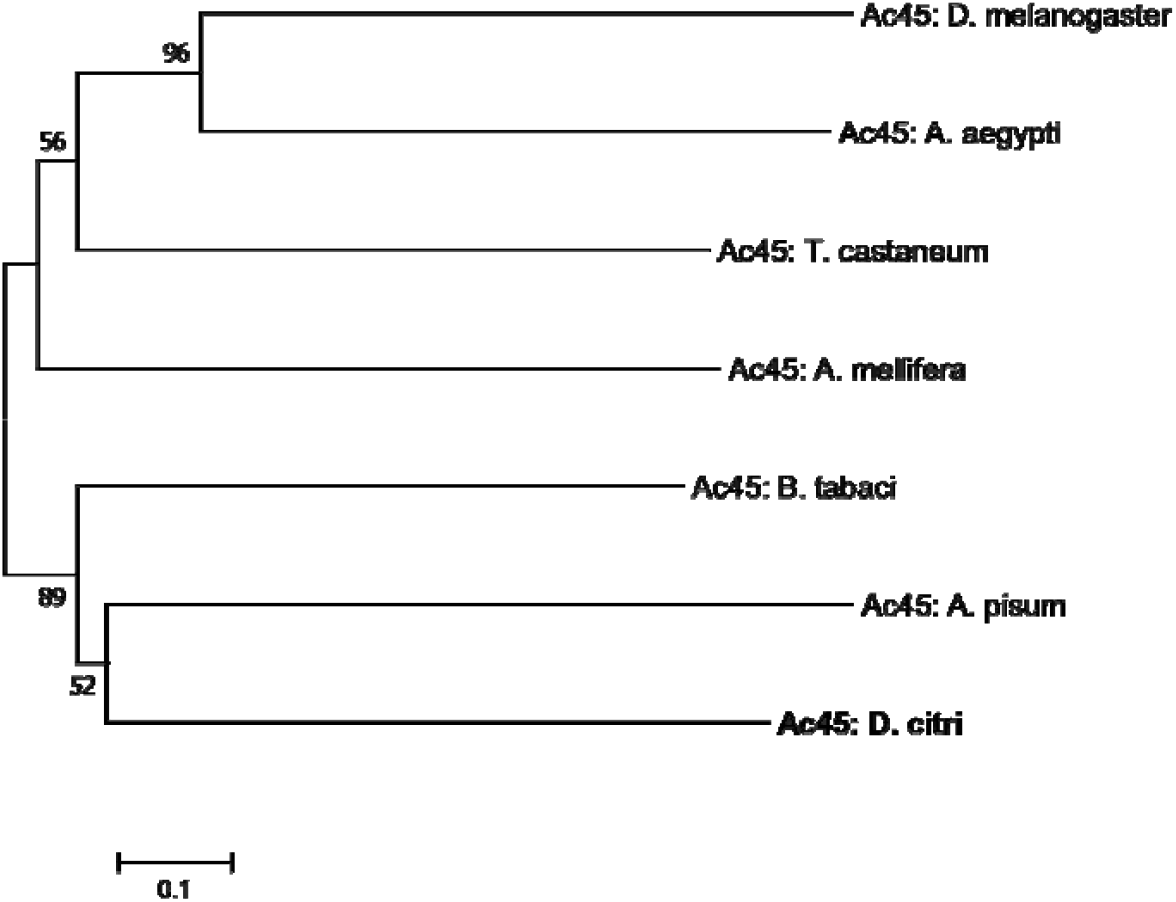
Phylogenetic analysis of V-ATPase accessory subunit Ac45. The tree was constructed with MEGA7 software [21] using MUSCLE for alignment of amino acid sequences, followed by neighbor-joining analysis with 1000 bootstrap replications. Values greater than 50 are shown at nodes. *D. citri is* marked in bold. NCBI accession numbers are shown in Table 4.

**Figure 5.**
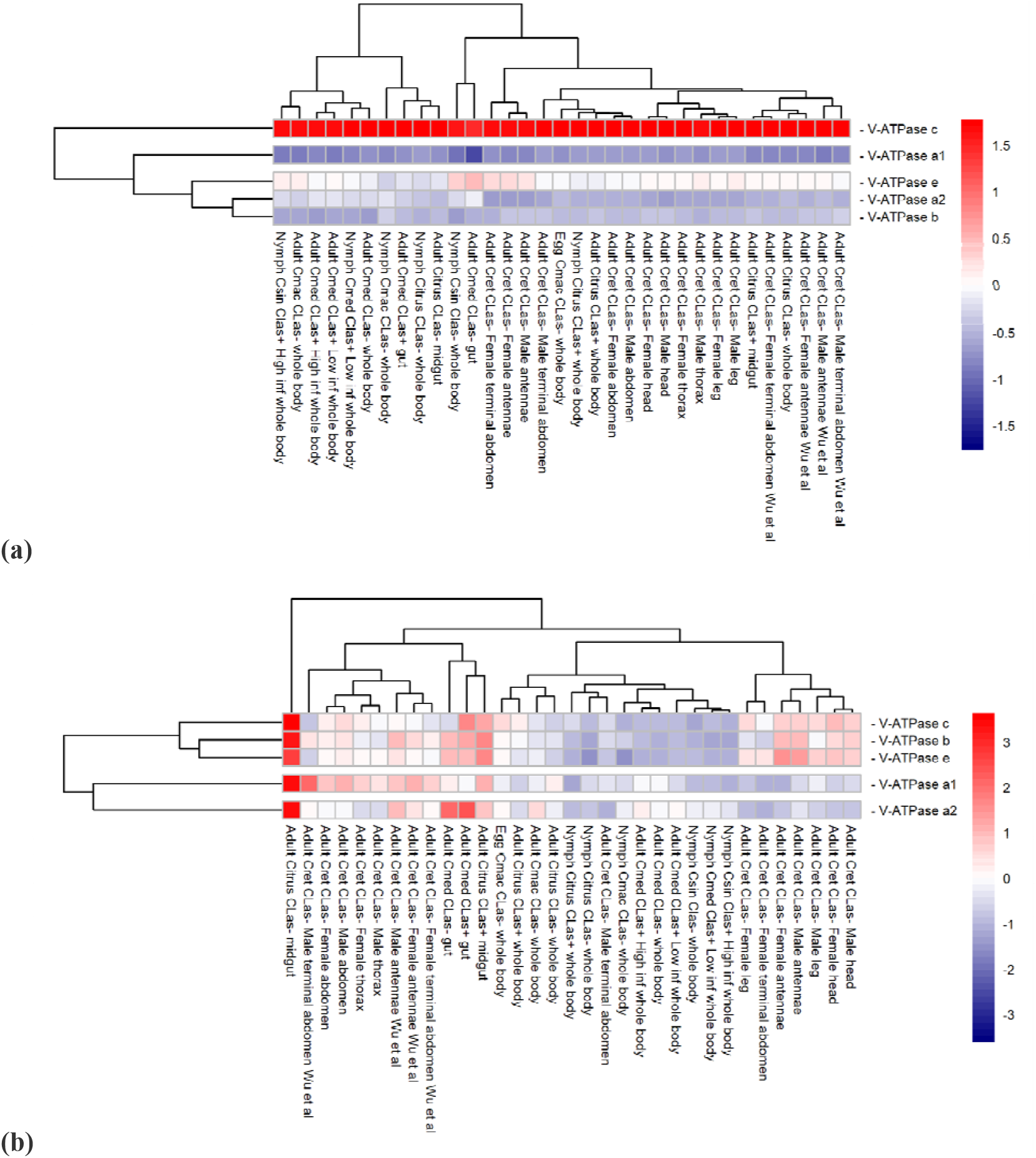
Comparative expression levels of the *D. citri V-ATPase* genes encoding the V_0_, transmembrane, subunits in *D. citri* insects reared on various infected and uninfected citrus varieties. Expression data were collected from the Citrus Greening Expression Network [9], with psyllid RNA-Seq data from NCBI BioProject’s PRJNA609978 and PRJNA448935, in addition to several published datasets [24–28]. Citrus hosts are abbreviated as Csin (*Citrus sinensis*), Cmed (*Citrus medica*), Cret (*Citrus reticulata*), and Cmac (*Citrus macrophylla*). TPM values are listed in Table 5. Rows of genes and columns of RNA-Seq data are clustered based on expression differences. **(a)** Expression data scaled by sample. **(b)** Expression data scaled by gene.

**Figure 6.**
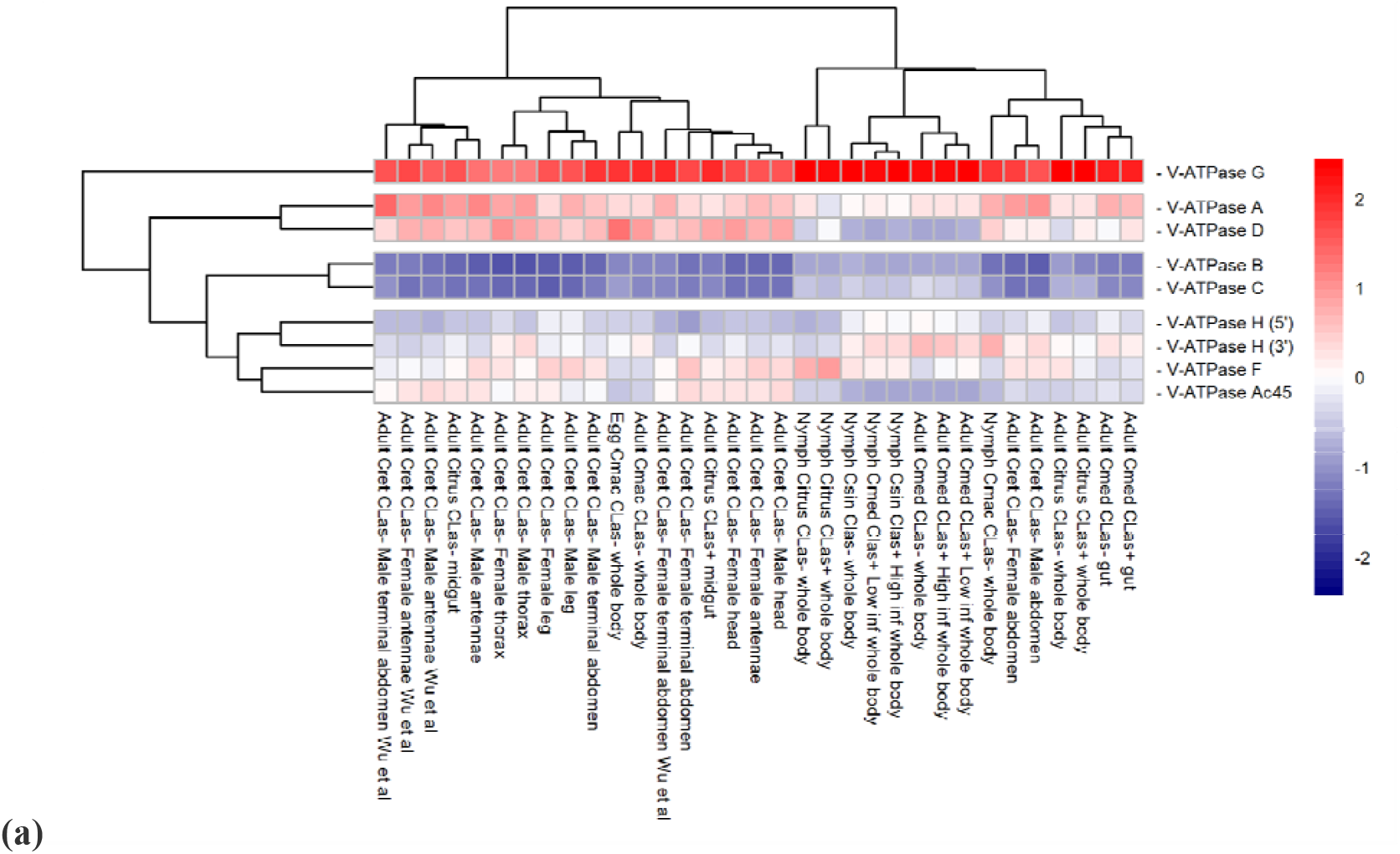
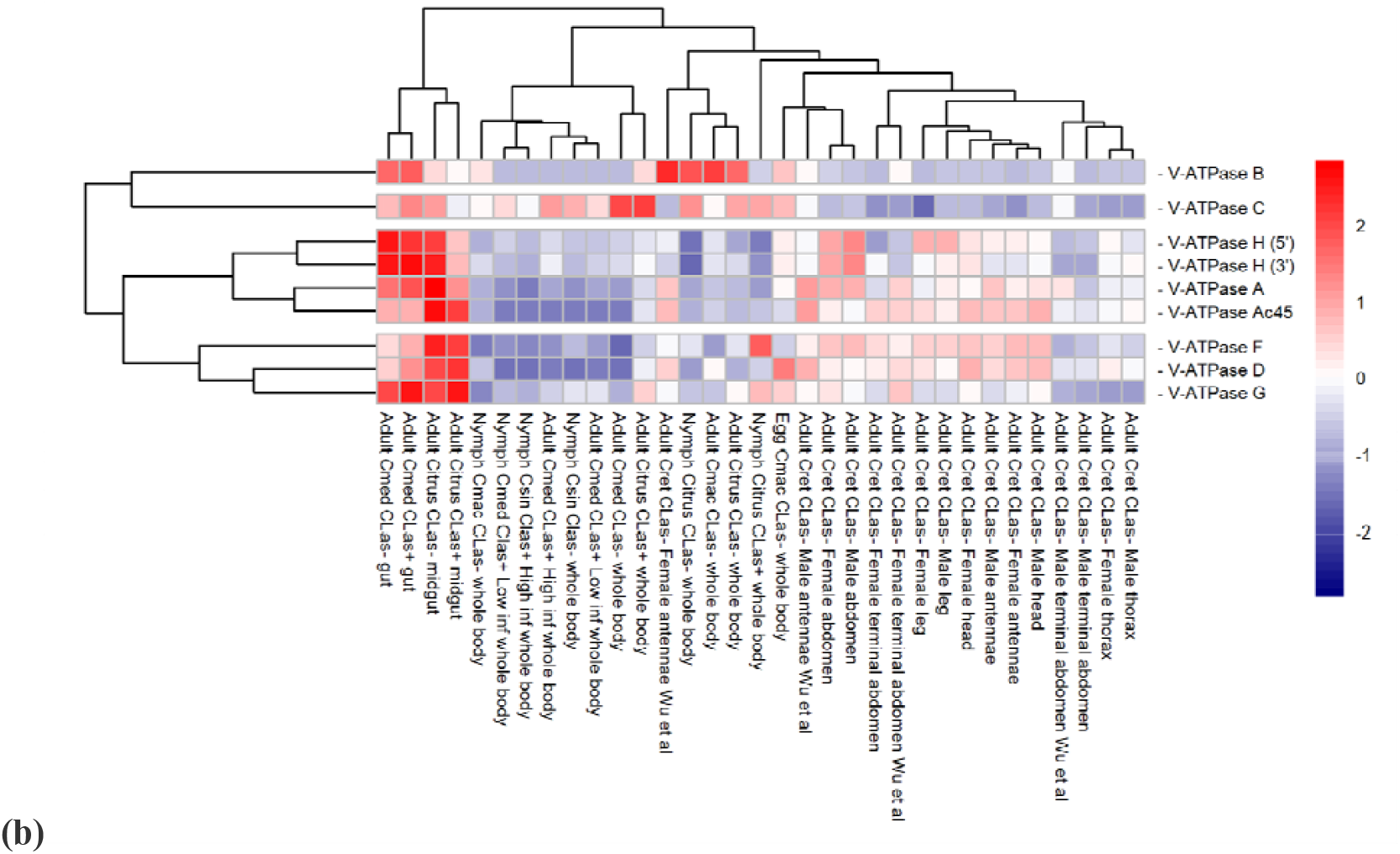
Comparative expression levels of the *D. citri V-ATPase* genes encoding the V_1_, catalytic, and accessory subunits in *D. citri* insects reared on various infected and uninfected citrus varieties. *V-ATPase H* was annotated as two partial models and are both represented here separately as *V-ATPase H* (5’), denoting the 5-prime end of the gene, and *V-ATPase H* (3’), denoting the 3-prime end of the gene. Expression data were collected from the Citrus Greening Expression Network [9], with psyllid RNA-Seq data from NCBI BioProject’s PRJNA609978 and PRJNA448935, in addition to several published datasets [24–28]. Citrus hosts are abbreviated as Csin (*Citrus sinensis*), Cmed (*Citrus medica*), Cret (*Citrus reticulata*), and Cmac (*Citrus macrophylla*). TPM values are listed in Table 5. Rows of genes and columns of RNA-Seq data are clustered based on expression differences. **(a)** Expression data scaled by sample. **(b)** Expression data scaled by gene.

### Data Validation and Quality Control

Genes encoding all 13 subunits required to build a single Vacuolar ATP synthase enzyme, as well as an accessory subunit S1 (*Ac45*) gene, were annotated in *D. citri*. There were no additional subunits found in *D. citri*, as reported in other metazoans [2]. Although insect V-ATPases are known to contain 13 subunits, there is variation in the gene copy number for individual subunits among different species (Tables 6-8). The V_0_ transmembrane domain subunits *V-ATPase a, b*, and *e*; the V_1_ catalytic domain subunits *V-ATPase A, C, D*, and *G*; and *Ac45*, all show variation in copy number among different species. The three Hemipterans analyzed (*D. citri, A. pisum*, and *B. tabaci*) maintain the same paralog number for all *V-ATPase* genes except for the *A. pisum V-ATPase D* and *G*, as compared to the other orders (Table 7). This variation in copy number is interesting in contrast to the genes *V-ATPase c, d, B, E, F*, and *H* that maintain only one gene copy across all Orders of Insecta found in Tables 6-8. V-ATPase subunits have been studied in plants, animals, fungi, and insects, and certain genes have been highlighted for their functional versatility in serving cell needs. For example, yeast and mammals have numerous copies and isoforms of the transmembrane proteolipid *V-ATPase a* with functions that support vacuoles, Golgi, neurons, osteoclasts, and epididymal cells [2]. *D. citri*, along with *A. pisum* and *B. tabaci*, has two copies of the *V-ATPase a* gene, whereas *D. melanogaster* has five copies (Table 6). In *D. citri*, a paralog of *V-ATPase a* was found, and they maintain differences in their amino acid sequences (Table 2, Table 6). Phylogenetic analysis of V_0_ subunit protein sequences supports that the duplication event occurred before the divergence of Hemimetabola and Holometabola (Figure 2).

**Table 6.**
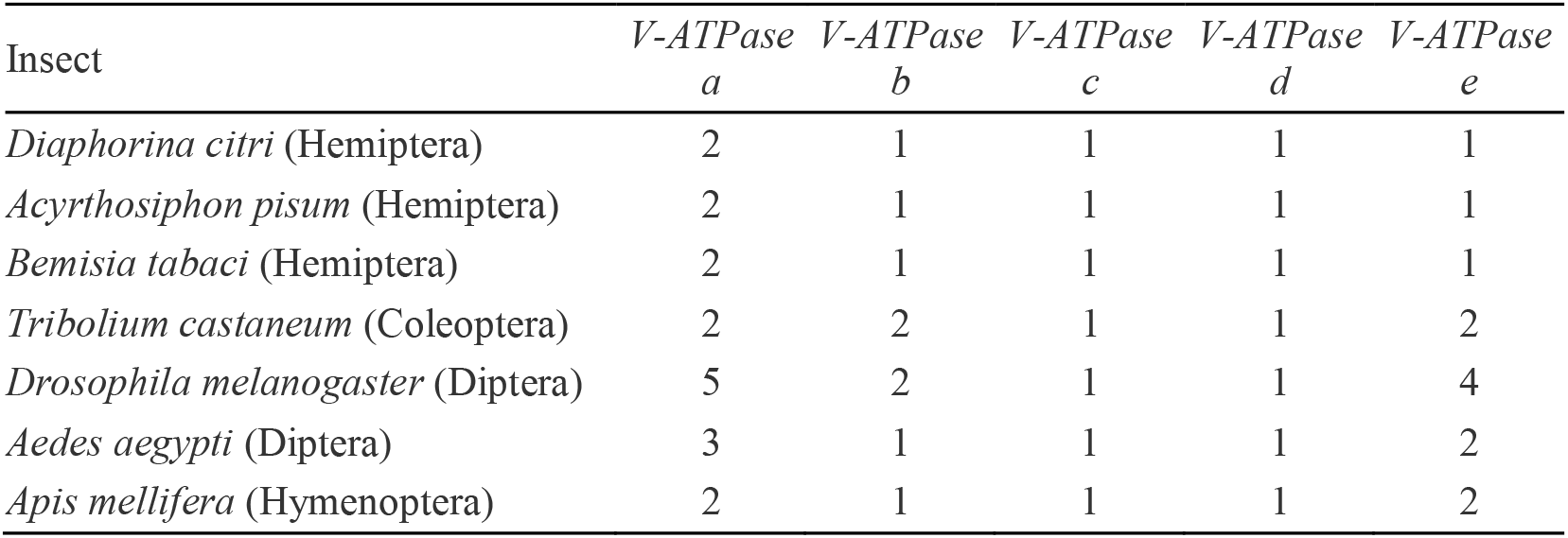
Gene copy comparison of V_0_ transmembrane subunit *V-ATPase* genes in *D. citri* and orthologous insect genes.

**Table 7.** Gene copy comparison of V_1_ catalytic subunit *V-ATPase* genes in *D. citri* and orthologous insect genes.

| Insect | *V-ATPase A* | *V-ATPase B* | *V-ATPase C* | *V-ATPase D* | *V-ATPase E* | *V-ATPase F* | *V-ATPase G* | *V-ATPase H* |
| --- | --- | --- | --- | --- | --- | --- | --- | --- |
| *Diaphorina citri* (Hemiptera) | 1 | 1 | 1 | 1 | 1 | 1 | 1 | 1 |
| *Acyrtosiphon pisum* (Hemiptera) | 1 | 1 | 1 | 2 | 1 | 1 | 2 | 1 |
| *Bemisia tabaci* (Hemiptera) | 1 | 1 | 1 | 1 | 1 | 1 | 1 | 1 |
| *Tribolium castaneum* (Coleoptera) | 1 | 1 | 1 | 3 | 1 | 1 | 2 | 1 |
| *Drosophila melanogaster* (Diptera) | 2 | 1 | 1 | 3 | 1 | 1 | 1 | 1 |
| *Aedes aegypti* (Diptera) | 1 | 1 | 2 | 2 | 1 | 1 | 3 | 1 |
| *Apis mellifera* (Hymenoptera) | 1 | 1 | 1 | 2 | 1 | 1 | 1 | 1 |

We were able to identify complete genes in genome v3.0 for all the subunits except *V-ATPase H*. Using genome-independent transcript sequences, we were able to determine that the 3’ portion of the *V-ATPase H* gene is located on chromosome 1, but the 5’ end of the gene is on one of the unplaced chromosomes that make up chromosome 0 (Table 1). Tables 2-4 show results of protein BLAST analysis comparing the same insects as found in Tables 6-8. Other than *Ac45*, all subunits share a relatively high identity, approximately 57-94%, among individual pairwise alignments with each *D. citri* sequence (Tables 2-3). BLAST results of annotated gene models had high query coverage to orthologs supporting the completeness of the annotated gene models. In contrast, the sequence identities of *Ac45*, approximately 24-33%, show the highest divergence when comparing *D. citri* to other insects (Table 4). For the V_0_, transmembrane domain, subunits in Table 2, proteolipid subunit c (*V-ATPase c*) maintains some of the highest percentages of sequence identity, highlighting the importance of the protein function to form the c-ring that rotates and ultimately translocates protons across various membranes [2]. This is supported in Table 6, in which a single gene copy for *V-ATPase c* is maintained across different orders of insects.

The Citrus Greening Expression Network (CGEN), found at citrusgreening.org, was used to compare transcript expression levels in various regions of *D. citri* which have either been exposed to or not exposed to *C*Las infection, the causative agent of citrus greening disease [9,29,30]. Figure 5 shows a heatmap comparing V_0_ subunit expression levels found under various conditions. *V-ATPase c* is visually differentiated by its inflated expression levels as compared to other *V-ATPase* transmembrane subunit genes (Figure 5a). *V-ATPase c* expression also shows a 2.63-fold increase, from 647.9 to 1705.25 TPM, in the guts of adult psyllids fed on infected *versus* uninfected *C. medica* leaves (Figure 5b, Table 5). These expression levels, coupled with the fundamental cellular nature and relatively even occurrence of *V-ATPases*, suggest that *V-ATPase* genes are good candidates for RNAi. Silencing a V_0_ transcript should have inhibitory effects on the assembly of the V-ATPase enzyme. In particular, if infected psyllids increase in their demand for higher *V-ATPase c* expression levels overall, knocking this transcript down will likely be detrimental for the insect. However, it cannot be determined at this time whether the elevated expression of *V-ATPase c* relative to other subunits in infected psyllids is due to higher demand of these proteins in the cell and should therefore be studied further in future research.

Of the V_1_, catalytic domain, subunit genes, *V-ATPase A* and *V-ATPase B* maintain the highest percentages of sequence identity, consistent with the importance of their function in containing the ATP binding sites at the V-ATPase subunits A/B protein interface (Table 3) [31]. Apart from *D. melanogaster, V-ATPase A* and *B* also maintain single copies of these two genes across different orders of insects, supporting their conserved nature as compared to other genes of this enzyme (Table 7). When comparing expression of the V_1_ subunits, *V-ATPase A* shows much higher expression compared to *V-ATPase B* across each measured variable, with *V-ATPase G* showing the highest expression in this group overall (Figure 6a). Unlike *V-ATPase c*, no significant differential expression was observed between the guts of insects reared on infected *versus* uninfected citrus trees (Figure 6b). However, *V-ATPase B* does show a reverse correlation, with a decrease in expression from 28.72 to 2.44 TPM in the whole body of *D. citri* nymphs raised on uninfected *versus* infected *Citrus spp*. (Table 5). A similar expression pattern can be seen throughout many of the *V-ATPase* catalytic genes and may infer an interaction between these genes and pathogen infection, therefore warranting further investigations (Table 5).

Figures 2, 3, and 4 depict phylogenetic analyses for the V_0_ transmembrane and V_1_ catalytic domains, and the Ac45 protein of V-ATPase, respectively. The individual V-ATPase subunits form clades, regardless of insect species. These clades also have the highest bootstrap values. This agrees with previous research that describes the enzyme as ancient and highly conserved. The evolution of V-ATPase has been analyzed for gene duplication and divergence from other ATP synthases, like F- and A-ATPase, which occur across the three domains of life [3]. Figures 2, 3, and 4 concur and suggest that the V-ATPase enzyme utilized in these insects existed in their common ancestor before they diverged into their respective species. The proteolipid subunit c and subunit d have the shortest branch lengths in Figure 2, consistent with Tables 2 and 6 which depict this to be of the most conserved subunits. Subunit c, which is required to form the critical c-ring rotor of V-ATPase [2], and subunit d, which may play a role as part of the central rotor of the V-ATPase [31], appear to have diverged the least when compared to the other transmembrane domain subunits and other insect species. In contrast, subunit e has diverged the most (Figure 3). This is consistent with the variable gene copy number observed across different orders of insects and the lower percentages of protein sequence identity seen in *D. citri* pairwise alignments (Tables 2,6). In addition, the function of subunit e is still unknown for the transmembrane domain subunits [5].

Figure 4 shows the evolutionary relatedness of the *D. citri* Ac45 protein. It is a relatively new protein critically associated with the assembly of a certain cell type V-ATPase and is still being studied [6]. For this select group of insect species, Ac45 groups and forms a clade with the other hemipteran protein sequences (Figure 4). *Ac45* is a variable gene when comparing *V-ATPase* across the domains of life, a paralog variability that is also seen among different orders of insects (Table 8) [6,32]. *Ac45* has diverged most of all the V-ATPase subunits in *D. citri* compared to other insects. This divergence is seen in phylogenetic analysis, denoted with longer branch lengths (Figure 5), and is also supported in the values of the pairwise alignments, in which the protein shares very little sequence identity across the query lengths (Table 4). Perhaps it is experimentally beneficial that the Ac45 protein shows the least conservation with other insect orthologs. It may serve as a species-specific targeted approach to limiting the psyllid from vectoring the causative agent of citrus greening disease while leaving related species unharmed and their ecology intact. However, Ac45 shows a markedly depressed transcription level as compared to other subunits (Figure 6a, Table 5). This likely reflects the limits in resolution with current whole RNA isolation and sequencing methods but still indicates the relatively low total expression, nonetheless. The Ac45 protein has not been observed to exist in every cell type depending on the organism and so is not necessarily utilized by every V-ATPase in the psyllid [6]. Thus, the expression data agree with previously published research.

**Table 8.** Gene copy comparison of the *Ac45* gene in *D. citri* and orthologous insect genes.

| Insect | *Ac45* |
| --- | --- |
| *Diaphorina citri* (Hemiptera) | 1 |
| *Acyrtosiphon pisum* (Hemiptera) | 1 |
| *Bemisia tabaci* (Hemiptera) | 1 |
| *Tribolium castaneum* (Coleoptera) | 2 |
| *Drosophila melanogaster* (Diptera) | 2 |
| *Aedes aegypti* (Diptera) | 2 |
| *Apis mellifera* (Hymenoptera) | 1 |

## Conclusion

The V-ATPase is a fundamental enzyme that functions exclusively as ATP-dependent proton pumps in almost every eukaryotic cell. V-ATPase allows for the proper functioning of endosomes and the Golgi apparatus, and it generates a proton-motive force in organelles and across plasma membranes that is utilized as a driving force for secondary transport processes [1]. Identification of these enzymes in the hemipteran, *D. citri*, provides a novel insect lineage for studies of insect evolution and biology, and may also provide potential targets for *D. citri*-specific molecular mechanisms for the management of HLB in citrus production systems [33– 35]. *D. citri* shows no deviation in the expected copy numbers of each of the *V-ATPase* genes (Tables 6-8). The data collected from *D. citri* reveals consistency among the genes previously characterized to be highly conserved, such as *V-ATPase c, d, A*, and *B* (Tables 2-4) [3,31]. While expression data was not available for *V-ATPase d, V-ATPase c* shows comparatively high expression levels overall and shows differential expression, 647.9 *versus* 1705.25 TPM, in the guts of adult psyllids fed on uninfected *versus* infected *C. medica* leaves (Figure 5, Table 5). Conversely, the *Ac45* gene shows low expression throughout life stages and tissues compared to other *V-ATPase* genes, however, the highly divergent nature of this gene may serve as a species-specific targeted approach to psyllid control (Table 4, Figure 6).

In hemipterans, RNAi efficacy has been successfully demonstrated for psyllids, whitefly, and leafhoppers [33–40]; planthoppers [41,42]; bedbugs [43]; and others [44–48]. RNAi targeting specifically the *V-ATPases* in hemipteran insects have been reported for the corn planthopper, *Peregrinus maidis* (Ashmead) (Hemiptera: Delphacidae) [12]; the corn leafhopper, *Dalbulus maidis* (Hemiptera: Cicadellidae) [13]; the Brown planthopper, *Nilaparvata lugens* (Stål) (Hemiptera: Delphacidae) [41]; and the bedbug, *Cimex lectularius* L. (Hemiptera: Cimicidae) [43], resulting in increased mortality and reduced fecundity. Thus, the highly divergent nature of these gene sequences provides unique targets that may serve as species-specific targeting for RNAi approaches in the management of psyllid vectors and other hemipteran pests [49,50].

### Reuse Potential

The manually curated gene models generated through this *D. citri* community annotation project will be available as part of the Official Gene Set version 3. Analysis of this data, including BLAST and expression profiling, can be conducted using the citrusgreenin.org website and Citrus Greening Expression Network (CGEN). The improved annotations presented in this study will facilitate experimental design to investigate the potential of *V-ATPases* as gene targets for therapies to control *D. citri*. Research considering differential expression patterns on V-ATPase transcripts in psyllids fed on *C*Las infected plants should be conducted. Additional studies are also required to confirm the role of the Ac45 protein, as its divergent nature may provide novel and species-specific gene targets, potentially through the use of RNAi, to control psyllid populations and reduce the effects of pathogens such as *C*Las.

## Declarations

## List of Abbreviations

ACP: Asian citrus psyllid
V-ATPase: Vacuolar (H+)-ATP synthase
V_0_: V-ATPase noncatalytic transmembrane domain
V_1_: V-ATPase catalytic cytoplasmic domain
ATP: Adenosine triphosphate
ADP: Adenosine diphosphate
RNAi: RNA interference
HLB: Huanglongbing
*C*Las: *Candidatus* Liberibacter asiaticus
HGNC: HUGO Gene Nomenclature Committee
NCBI: National Center for Biotechnology Information
MCOT: Maker, Cufflinks, Oasis, Trinity
BLASTp: protein BLAST
RNA-seq: RNA sequencing
DNA-seq: DNA sequencing
Iso-seq: Isoform sequencing
OGS: Official Gene Set
ChrXX: Chromosome number location of OGS
QC: Query coverage
CGEN: Citrus Greening Expression Network
TPM: Transcripts per million
Csin: *Citrus sinensis*
Cmed: *Citrus medica*
Cret: *Citrus reticulate*
Cmac: *Citrus macrophylla*

## Ethical Approval

Not applicable.

## Consent for publication

Not applicable.

## Authors’ contributions

WBH, SJB, TD, and LAM conceptualized the study; TD, SS, TDS, and SJB supervised the study; SJB, TD, SS, and LAM contributed to project administration; RG conducted investigation; PH, MF-G, and SS contributed to software development; PH, MF-G, SS, TDS, and JB developed methodology; SJB, TD, WBH, and LAM acquired funding; RG and CM prepared and wrote the original draft; TD, SJB, SS, TDS, WT, WBH and JB reviewed and edited the draft.

## Acknowledgments

We would like to thank Helen Wiersma-Koch (Indian River State College) for her assistance.

